# Phylogenetic profiling identifies glucosyl-phosphoryl dolichol scramblase candidates

**DOI:** 10.1101/209106

**Authors:** Lucy A. Skrabanek, Anant K. Menon

## 1 Background

Protein glycosylation is essential for all eukaryotes, from disease-causing protists such as malaria and trypanosomes, to yeast and mammals. Secretory proteins are almost invariably *N*-glycosylated, *O*- and *C*-mannosylated, and/or GPI-anchored as they enter the lumen of the endoplasmic reticulum (ER).

All ER protein glycosylation reactions occur in the *lumen* and often involve lumenal mannosylation and glucosylation steps in which mannose and glucose residues are sourced from the glycolipids mannosyl- and glucosyl-phosphoryl dolichol (MPD and GPD, respectively) [1, 2]. Paradoxically, these two lipids are synthesized on the *cytoplasmic* face of the ER and must therefore be flipped across the ER membrane to provide a source of *lumenal* mannose and glucose. As the spontaneous rate of MPD and GPD flipping is extremely low, specific transporters are needed to facilitate the transbilayer movement of MPD and GPD across the ER membrane at a physiological rate. MPD and GPD transport activities have been demonstrated and characterized in ER microsomes, as well as in vesicles reconstituted with ER membrane proteins [3–6]. The transport proteins have been shown to be highly structure specific, discriminating between isomers of their lipid substrates, and facilitating lipid movement bidirectionally in an ATP-independent manner (the last point defines them as *scramblases*, whereas they were previously known as ATP-independent flippases). Although most of the enzymes and co-factors of ER protein glycosylation are known, the molecular identities of the critical dolichol glycolipid scramblases remain a mystery.

Unlike MPD scramblase that is required for all ER protein glycosylation reactions, GPD scramblase is needed exclusively for the synthesis of the glucosylated *N*-glycan precursor Glc_3_Man_9_GlcNAc_2_-PP-dolichol (G3M9-DLO). Even though non-glucosylated *N*-glycan precursors are substrates for the protein *N*-glycosylation machinery, the presence of the tri-glucosyl cap – and hence GPD scramblase activity – is critically important for glycosylation efficiency in many eukaryotes, including yeast and humans [7]. Two points are noteworthy. (i) Not all organisms have glucose in their *N*-glycan precursor [8]. (ii) While the synthesis of glucosylated *N*-glycan precursors is not essential for the viability of yeast [9, 10], yeast cells that are deficient in G3M9-DLO synthesis display numerous phenotypes including under-glycosylation of proteins, abnormal cell shape and altered susceptibility to a variety of chemicals. In humans, optimization of oligosaccharide transfer efficiency provided by the glucosyl cap is critical as evinced by severe human diseases.

Taking advantage of the fact that not all *N*-glycosylation-competent organisms have glucose in their *N*-glycan precursor, we implemented a bioinformatics approach for assignment of protein function, namely phylogenetic profiling. Using this procedure, we identified a number of polytopic ER membrane proteins as GPD scramblase candidates in yeast.

## 2 Results

### 2.1 GPD scramblase candidates identified by phylogenetic profiling

Phylogenetic profiling predicts protein function based on patterns of protein presence or absence across multiple species (e.g. [11, 12]). If homologs are inherited or lost co-dependently, there is a high chance that they are functionally related or physically interacting, because they are likely to be subject to the same functional constraints, or lack thereof. Proteins with similar phylogenetic profiles tend to be part of a functional unit. A previous study [13] showed that glycosyltransferases are present or absent in organisms in sets. Here we used patterns generated by phylogenetic profiling to identify GPD scramblase candidates. As not all organisms have glucose in their *N*-glycan precursor [8] we hypothesized that the presence or absence of GPD scramblase in a particular organism will be highly correlated with, but not necessarily identical to, the presence or absence of other proteins of the glucosylation pathway, e.g. the glucosyltransferase Alg6 (Figure 1). We used yeast as a reference organism as it has a well-annotated genome and a complete *N*-glycosylation pathway.

**Figure 1:**
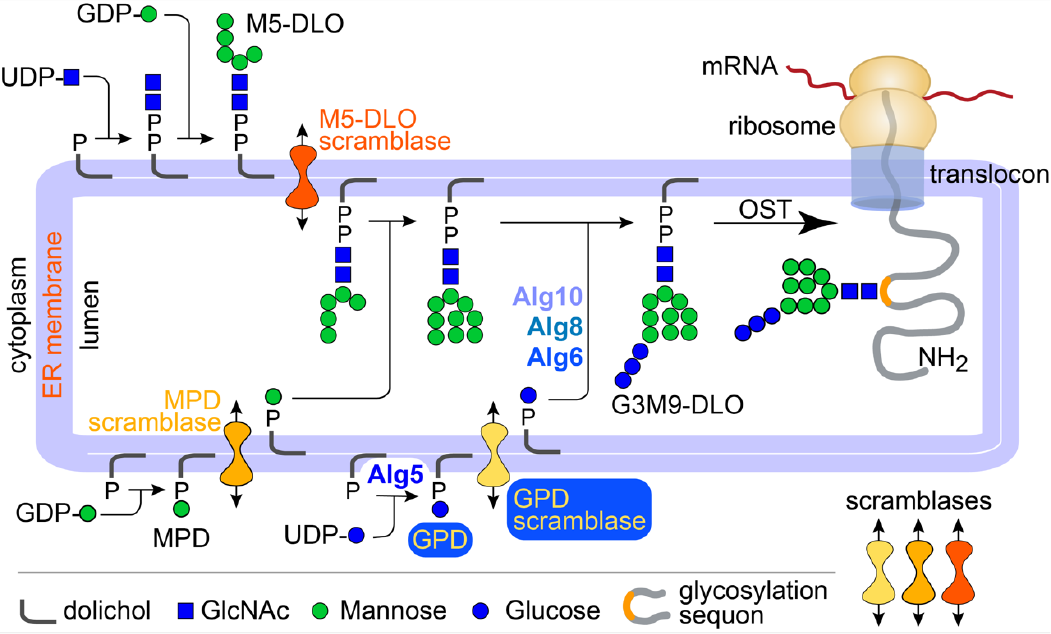
Role of GPD scramblase in G3M9-DLO synthesis in yeast. Glc_3_Man_9_GlcNAc_2_-PP-dolichol (G3M9-DLO; DLO=dolichol-linked oligosaccharide) is synthesized via a series of reactions that begin on the cytoplasmic face of the ER to generate Man_5_GlcNAc_2-_PP-dolichol (M5-DLO). M5-DLO is translocated across the ER membrane by M5-DLO scramblase. In the lumen, M5-DLO is extended to Man_9_GlcNAc_2_-PP-dolichol (M9-DLO) through the action of MPD-dependent mannosyltransferases and eventually extended to G3M9-DLO. The glucosylation reactions require lumenally oriented GPD. GPD is synthesized from dolichyl-P and UDP-glucose on the cytoplasmic face of the ER by GPD synthase (Alg5), and then moved to the lumen by GPD scramblase. In the lumen, the glucosyltransferases Alg6, Alg8 and Alg10 use GPD to add glucose residues to M9-DLO to generate G3M9-DLO that provides the oligosaccharide used by oligosaccharyltransferase (OST) to *N*-glycosylate proteins. The synthesis, scrambling and consumption of GPD likely form a co-evolving metabolic unit. This premise is the basis of our phylogenetic profiling identification of GPD scram-blase candidates.

### 2.2 Annotated genomes for phylogenetic profiling

We downloaded 687 annotated genomes from the Ensembl database: 408 fungal, 133 protist, 143 archaeal, 3 higher eukaryotes (dog, mouse, human). We downloaded all the fungal and all the protist genomes from release 27 of the Ensembl database, as well as 143 selected archaeal genomes (those which are also part of the OMA Browser database).

Within this set we identified organisms capable of *N*-glycosylation by using BLAST [14] (E-value threshold 1e-6) to probe for Alg7, the enzyme that initiates DLO synthesis, and Stt3, the catalytic subunit of oligosaccharyltransferase (we identified the Alg7 and Stt3 proteins in Saccharomyces cerevisiae from the SwissProt database as entries P07286.1 and P39007.2, respectively, which correspond to the YBR243C and YGL022W proteins in the Ensembl (SGD) annotation). Only genomes encoding both proteins were retained.

Phylogenetic profiling benefits from an increased amount of input data, but previous studies have shown diminishing returns as greater numbers of genomes are added [15]. Thus, we further reduced our dataset by picking a single exemplar for species that were represented by more than one strain. Our final list contained 337 genomes (see Appendix A).

### 2.3 Species tree of organisms capable of *N*-glycosylation

Several authors have suggested that phylogenetic profiles should be ordered by the phylogenetic relatedness of the constituent organisms, e.g., Cokus et al. [16]. The 337 organisms being used in this analysis were imported into NCBI’s Taxonomy Browser (http://www.ncbi.nlm.nih.gov/Taxonomy/CommonTree/wwwcmt.cgi). Some species were represented in NCBI under a different name, either because the name was spelled slightly differently in NCBI, or because the asexual and sexual stages have different names; these were accordingly changed before import. Table 1 indicates the species which are named differently in the Ensembl and NCBI databases.

**Table 1:**
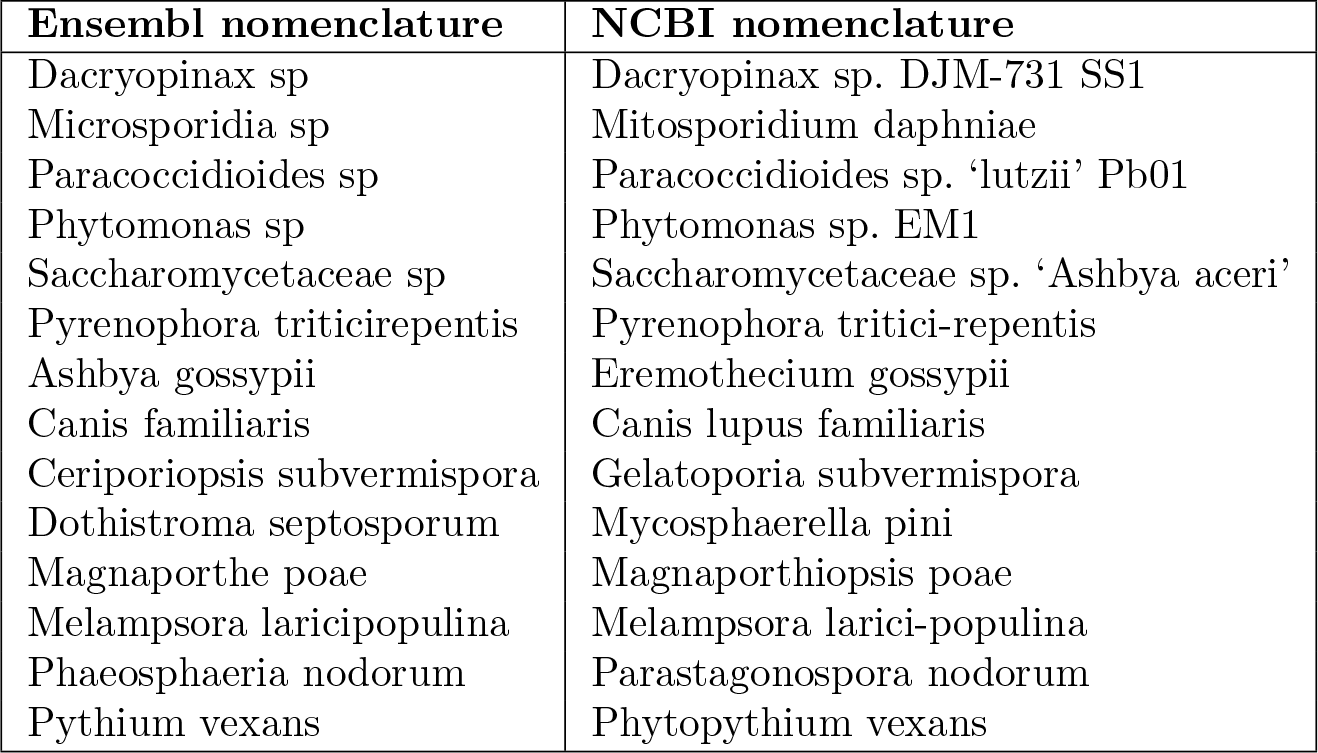
Species whose names differ in the Ensembl and NCBI databases.

The phylogenetic representation of the relationship of each of the 337 organisms to each other as determined by the NCBI’s Taxonomic Browser was retrieved. The order of the organisms shown in Appendix A reflects the phylogenetic relationship to one another as determined by this method.

### 2.4 Profile generation

We used BLAST [14] to search the protein complement of each of the 337 genomes for homologs to all 6,692 yeast proteins. This was done three separate times, each time using a different E-value threshold cutoff (no threshold, 1e-2, and 1e-10 for our most stringent match). Three types of profiles were generated: (i) **BRH** (best reciprocal hit), a binary profile, where the presence or absence of each yeast protein is predicted for each organism. For a yeast protein to be marked as existing in organism O, the protein in organism O that is the highest scoring hit to the yeast protein must also find that same yeast protein as one of its top two hits when searched against the yeast proteome. (ii) **Score profile**, a quantitative homology measure, where the score of the top hit to the yeast protein is divided by the score of its self-match (i.e., the score when the yeast protein is aligned to itself). This normalized score is a continuous variable from 0 to 1 (0 = non-existence; 1 = perfect match). (iii) **E-value profile**, calculated by taking the 1/log10(E-value score), and capping the maximum value at 1. The E-value profile is on a 0-1 scale, where 0 is a perfect match, and 1 is non-existence. The profile of Alg5 (GPD synthase) [9, 10] was more promiscuous than expected (see below), so as bait we instead used the GPD-dependent glucosyltransferase Alg6 [9]. Distances of all profiles were calculated relative to Alg6, using a Jaccard-like metric for the BRH profile, and the Canberra distance metric for the others.

### 2.5 Filters

Annotations for each yeast protein were obtained from the Saccharomyces Genome Database (SGD) (http://downloads.yeastgenome.org/curation/chromosomal_feature/SGD_features.tab). Only records pertaining to ORFs were retained. Transmembrane domains were predicted for all yeast proteins using the TMHMM server (http://www.cbs.dtu.dk/services/TMHMM/). [17]). The results were filtered to include only membrane proteins, and further refined to prioritize only multispanning proteins (≥3 transmembrane domains). We also filtered out proteins that were not found in the three higher eukaryotes. A final manual filter restricted the list to proteins that are known to be ER-localized, and those whose localization is ambiguous.

For each BLAST E-value threshold, we listed the top 50 hits that were identified in all three profile-generating methods (Appendix B) and ranked them according to the lowest rank that a protein has in any of the three lists. Although the lists differ slightly, the top candidates are common to all: (in rank order) Alg8, Mns1, Ale1, Erg24, Erc1, Ybr220C, Erg3, Gwt1, Alg2, Scs7, Ste24, Erg11, Sur2, Ydr338C, Alg3, Alg9, and Ykr051W. Depending on the outcome of our biochemical analyses, we may pick additional candidates from the ranked lists that we generated, potentially including proteins that span the membrane only once. Such proteins may oligomerize to generate a pseudo-polytopic membrane protein that could have a transport function.

The presence of Alg8 as our top hit is unsurprising because it is required for synthesis of the glucose cap in G3M9-DLO and would be expected to co-evolve with Alg6. Three candidates (Ybr220C, Ydr338C, Ykr051W) are annotated as proteins of unknown function, two (Erc1, Ydr338C) are homologous to proteins that belong to the MOP exporter superfamily, and all except three (Gwt1, Alg2, Erg11) are non-essential for growth, consistent with glucosylation being non-essential in yeast. Interestingly, most have a lipid-related function, e.g. Ale1 is a lysophospholipid acyltransferase and Scs7 is a sphingolipid fatty acid α-hydroxylase, and three (Erg24, Ste24, Erg11) were identified in photocrosslinking studies using dolichyl phosphate analogs [18].

### 2.6 Notes on the profiling method and results

Phylogenetic profiling is a powerful method to identify functionally associated, co-evolving proteins such as enzymes that catalyze sequential steps in a biochemical pathway. While the method identifies the most significant co-evolutionary events, the generated profiles may not always match exactly. The most common causes for imperfect matches are a) incomplete protein annotation of a genome, b) different rates of evolution in independent lineages which may cause evolutionarily more distant proteins to outscore (in BLAST) the proteins which are actually closer, and (c) the possibility that in some organisms, a protein may have another, unrelated, function [11, 19].

Thus, Alg5 is found in a few organisms that do not synthesize glucosylated DLOs, suggesting novel roles for GPD [20] and/or an alternate function for Alg5 in these organisms. Likewise, the glucosyltransferase Alg8 is present in every Alg6-positive genome (268 of the 337 genomes we analyzed are Alg6-positive) as expected, but it is also present in 3 genomes (e.g. *Toxoplasma gondii*) that do not have Alg6. As Alg8 cannot add glucose unless Alg6 has first acted, and it has no other known function, its presence in these three genomes may be an evolutionary remnant. Another reason that profiles may not match exactly is if the GPD scramblase protein has another function, and is therefore subject to selective pressure unrelated to the glucosylation pathway. In this case, the candidate protein should be present if Alg6 is present, but may also be present when Alg6 is absent. For example, Erg24 is found in 19 of the 69 genomes that do not have Alg6. These genomes include those of trypanosomatids (e.g. *Trypanosoma brucei*) that do not synthesize glucosylated DLOs, but require the C14 sterol reductase activity of Erg24 to synthesize sterols. Thus, Erg24 may be a bifunctional protein with both GPD scramblase and sterol reductase activities. Despite the not-always-perfect profile matches, it is important to recognize that the power of the phylogenetic profiling approach used here lies in its ability to generate an unbiased, ranked list of candidates that allow us to prioritize our efforts to evaluate potential scramblase functions of the candidates.

## 3 Summary

Our top GPD scramblase candidates (in rank order, those in bold are essential for yeast growth) identified by phylogenetic profiling are Alg8, Mns1, Ale1, Erg24, Erc1, Ybr220C, Erg3, Gwt1, Alg2, Scs7, Ste24, Erg11, Sur2, Ydr338C, Alg3, Alg9, and Ykr051W. We are currently evaluating these candidates via *in vitro* and *in vivo* tests to identify the scramblase. This preprint will be updated once experimental data are available.

## Acknowledgements

We thank John Samuelson (Boston University Medical School) for his initial efforts on this project, and Sam Canis for stimulation. This work was supported by NIH grant NS093457 and a grant (P 61397-009) from the Alice Bohmfalk Charitable Trust (both to AKM).

## Appendix A: List of organisms used in phylogenetic profiling analysis, ordered by phylogenetic relatedness

1. Guillardia theta
2. Bigelowiella natans
3. Reticulomyxa filosa
4. Perkinsus marinus atcc 50983
5. Oxytricha trifallax gca 000295675
6. Stylonychia lemnae
7. Tetrahymena thermophila
8. Paramecium tetraurelia
9. Babesia equi strain wa
10. Babesia bovis
11. Plasmodium reichenowi
12. Plasmodium falciparum
13. Plasmodium inui san antonio 1
14. Plasmodium vivax
15. Plasmodium knowlesi
16. Plasmodium cynomolgi strain b
17. Plasmodium yoelii yoelii
18. Plasmodium vinckei petteri
19. Plasmodium chabaudi
20. Plasmodium berghei
21. Gregarina niphandrodes
22. Hammondia hammondi
23. Toxoplasma gondii
24. Cryptosporidium muris rn66
25. Cryptosporidium parvum iowa ii
26. Eimeria necatrix
27. Eimeria tenella gca 000499545
28. Naegleria gruberi
29. Polysphondylium pallidum pn500
30. Dictyostelium fasciculatum
31. Dictyostelium purpureum
32. Entamoeba nuttalli p19
33. Entamoeba dispar saw760
34. Entamoeba invadens ip1
35. Entamoeba histolytica
36. Acanthamoeba castellanii str neff
37. Spironucleus salmonicida
38. Giardia intestinalis
39. Giardia lamblia
40. Trichomonas vaginalis g3
41. Phytomonas sp isolate em1
42. Angomonas deanei
43. Trypanosoma rangeli sc58
44. Trypanosoma cruzi
45. Trypanosoma brucei
46. Leishmania mexicana mhom gt 2001 u1103
47. Leishmania major
48. Leishmania infantum jpcm5
49. Leishmania donovani
50. Leishmania panamensis
51. Leishmania braziliensis mhom br 75 m2904
52. Salpingoeca rosetta
53. Monosiga brevicollis mx1
54. Canis familiaris
55. Mus musculus
56. Homo sapiens
57. Rhizophagus irregularis daom 181602
58. Microsporidia sp ugp3
59. Nematocida parisii ertm1
60. Rozella allomycis csf55
61. Batrachochytrium dendrobatidis jam81
62. Wallemia ichthyophaga exf 994
63. Wallemia sebi cbs 633 66
64. Mixia osmundae iam 14324
65. Melampsora laricipopulina
66. Puccinia triticina
67. Puccinia graminis
68. Rhodosporidium toruloides np11
69. Microbotryum violaceum
70. Tilletiaria anomala ubc 951
71. Pseudozyma brasiliensis
72. Pseudozyma hubeiensis sy62
73. Pseudozyma aphidis dsm 70725
74. Pseudozyma antarctica
75. Sporisorium reilianum
76. Ustilago hordei
77. Ustilago maydis
78. Dacryopinax sp djm 731 ss1
79. Heterobasidion irregulare tc 32 1
80. Rhizoctonia solani 123e
81. Botryobasidium botryosum fd 172 ss1
82. Tulasnella calospora mut 4182
83. Gloeophyllum trabeum atcc 11539
84. Serendipita vermifera maff 305830
85. Piriformospora indica dsm 11827
86. Fibroporia radiculosa
87. Phanerochaete carnosa hhb 10118 sp
88. Phlebiopsis gigantea 11061 1 cr5 6
89. Ceriporiopsis subvermispora b
90. Fomitopsis pinicola fp 58527 ss1
91. Trametes cinnabarina
92. Jaapia argillacea mucl 33604
93. Piloderma croceum f 1598
94. Paxillus rubicundulus ve08 2h10
95. Serpula lacrymans var lacrymans s7 3
96. Scleroderma citrinum foug a
97. Pisolithus microcarpus 441
98. Pisolithus tinctorius marx 270
99. Suillus luteus uh slu lm8 n1
100. Moniliophthora roreri mca 2997
101. Galerina marginata cbs 339 88
102. Hebeloma cylindrosporum h7
103. Amanita muscaria koide bx008
104. Laccaria amethystina laam 08 1
105. Laccaria bicolor s238n h82
106. Coprinopsis cinerea okayama7 130
107. Agaricus bisporus var bisporus h97
108. Schizophyllum commune h4 8
109. Pleurotus ostreatus pc15
110. Trichosporon asahii var asahii cbs 2479
111. Cryptococcus gattii r265
112. Cryptococcus neoformans
113. Tuber melanosporum
114. Dactylellina haptotyla cbs 200 50
115. Arthrobotrys oligospora atcc 24927
116. Pseudogymnoascus destructans 20631 21
117. Pseudogymnoascus pannorum vkm f 103
118. Oidiodendron maius zn
119. Marssonina brunnea f sp multigermtubi mb m1
120. Glarea lozoyensis atcc 20868
121. Sclerotinia borealis f 4157
122. Botrytis cinerea
123. Erysiphe necator
124. Blumeria graminis
125. Pestalotiopsis fici w106 1
126. Eutypa lata ucrel1
127. Togninia minima ucrpa7
128. Grosmannia clavigera kw1407
129. Ophiostoma piceae uamh 11346
130. Sporothrix brasiliensis 5110
131. Sporothrix schenckii atcc 58251
132. Magnaporthe oryzae
133. Magnaporthe poae
134. Gaeumannomyces graminis
135. Myceliophthora thermophila atcc 42464
136. Chaetomium thermophilum var thermophilum dsm 1495
137. Chaetomium globosum cbs 148 51
138. Thielavia terrestris nrrl 8126
139. Podospora anserina s mat
140. Sordaria macrospora
141. Neurospora tetrasperma fgsc 2508
142. Neurospora crassa
143. Scedosporium apiospermum
144. Verticillium alfalfae vams 102
145. Verticillium dahliae
146. Colletotrichum sublineola
147. Colletotrichum fioriniae pj7
148. Colletotrichum gloeosporioides
149. Colletotrichum higginsianum
150. Colletotrichum graminicola
151. Colletotrichum orbiculare
152. Stachybotrys chlorohalonata ibt 40285
153. Stachybotrys chartarum ibt 40288
154. Beauveria bassiana arsef 2860
155. Cordyceps militaris cm01
156. Trichoderma atroviride imi 206040
157. Trichoderma reesei
158. Trichoderma virens
159. Fusarium solani
160. Fusarium pseudograminearum
161. Fusarium graminearum
162. Fusarium oxysporum
163. Fusarium verticillioides
164. Fusarium fujikuroi
165. Torrubiella hemipterigena
166. Metarhizium majus arsef 297
167. Metarhizium robertsii
168. Metarhizium brunneum arsef 3297
169. Metarhizium guizhouense arsef 977
170. Metarhizium acridum cqma 102
171. Metarhizium album arsef 1941
172. Metarhizium anisopliae
173. Claviceps purpurea 20 1
174. Ustilaginoidea virens
175. Acremonium chrysogenum atcc 11550
176. Endocarpon pusillum z07020
177. Cyphellophora europaea cbs 101466
178. Rhinocladiella mackenziei cbs 650 93
179. Cladophialophora psammophila cbs 110553
180. Cladophialophora immunda
181. Cladophialophora yegresii cbs 114405
182. Cladophialophora bantiana cbs 173 52
183. Cladophialophora carrionii cbs 160 54
184. Coniosporium apollinis cbs 100218
185. Fonsecaea pedrosoi cbs 271 37
186. Exophiala aquamarina cbs 119918
187. Exophiala sideris
188. Exophiala xenobiotica
189. Exophiala oligosperma
190. Exophiala mesophila
191. Exophiala spinifera
192. Exophiala dermatitidis nih ut8656
193. Capronia coronata cbs 617 96
194. Capronia epimyces cbs 606 96
195. Capronia semiimmersa
196. Byssochlamys spectabilis no 5
197. Talaromyces marneffei atcc 18224
198. Talaromyces stipitatus atcc 10500
199. Neosartorya fischeri
200. Penicillium rubens wisconsin 54 1255
201. Penicillium oxalicum 114 2
202. Penicillium solitum
203. Penicillium italicum
204. Penicillium digitatum pd1
205. Penicillium expansum
206. Aspergillus fumigatus
207. Aspergillus ruber cbs 135680
208. Aspergillus nidulans
209. Aspergillus terreus
210. Aspergillus oryzae
211. Aspergillus niger
212. Aspergillus flavus
213. Aspergillus clavatus
214. Coccidioides posadasii str silveira
215. Paracoccidioides sp lutzii pb01
216. Paracoccidioides brasiliensis pb03
217. Uncinocarpus reesii 1704
218. Microsporum gypseum cbs 118893
219. Arthroderma otae cbs 113480
220. Arthroderma benhamiae cbs 112371
221. Trichophyton soudanense cbs 452 61
222. Trichophyton equinum cbs 127 97
223. Trichophyton verrucosum hki 0517
224. Trichophyton interdigitale h6
225. Trichophyton tonsurans cbs 112818
226. Trichophyton rubrum d6
227. Blastomyces dermatitidis atcc 18188
228. Histoplasma capsulatum h88
229. Verruconis gallopava
230. Neofusicoccum parvum ucrnp2
231. Macrophomina phaseolina ms6
232. Baudoinia compniacensis uamh 10762
233. Zymoseptoria tritici
234. Sphaerulina musiva so2202
235. Pseudocercospora fijiensis cirad86
236. Dothistroma septosporum
237. Aureobasidium subglaciale exf 2481
238. Aureobasidium melanogenum cbs 110374
239. Aureobasidium pullulans exf 150
240. Phaeosphaeria nodorum
241. Leptosphaeria maculans
242. Setosphaeria turcica et28a
243. Pyrenophora teres
244. Pyrenophora triticirepentis
245. Bipolaris oryzae atcc 44560
246. Bipolaris sorokiniana nd90pr
247. Bipolaris victoriae fi3
248. Bipolaris zeicola 26 r 13
249. Bipolaris maydis atcc 48331
250. Kuraishia capsulata cbs 1993
251. Clavispora lusitaniae atcc 42720
252. Yarrowia lipolytica
253. Saccharomyces arboricola h 6
254. Saccharomycetaceae sp ashbya aceri
255. Kazachstania naganishii cbs 8797
256. Kazachstania africana cbs 2517
257. Lachancea lanzarotensis
258. Lachancea thermotolerans cbs 6340
259. Tetrapisispora blattae cbs 6284
260. Tetrapisispora phaffii cbs 4417
261. Vanderwaltozyma polyspora dsm 70294
262. Eremothecium cymbalariae dbvpg 7215
263. Ashbya gossypii
264. Kluyveromyces lactis
265. Naumovozyma dairenensis cbs 421
266. Naumovozyma castellii cbs 4309
267. Candida glabrata
268. Zygosaccharomyces rouxii
269. Zygosaccharomyces bailii isa1307
270. Torulaspora delbrueckii
271. Wickerhamomyces ciferrii
272. Komagataella pastoris
273. Spathaspora passalidarum nrrl y 27907
274. Candida tenuis atcc 10573
275. Lodderomyces elongisporus nrrl yb 4239
276. Candida orthopsilosis co 90 125
277. Candida dubliniensis cd36
278. Candida tropicalis mya 3404
279. Candida albicans wo 1
280. Debaryomyces hansenii cbs767
281. Meyerozyma guilliermondii atcc 6260
282. Scheffersomyces stipitis cbs 6054
283. Millerozyma farinosa cbs 7064
284. Ogataea parapolymorpha dl 1
285. Pichia kudriavzevii
286. Pneumocystis murina b123
287. Schizosaccharomyces cryophilus
288. Schizosaccharomyces octosporus
289. Schizosaccharomyces japonicus
290. Schizosaccharomyces pombe
291. Aureococcus anophagefferens
292. Blastocystis hominis
293. Albugo laibachii
294. Aphanomyces invadans
295. Aphanomyces astaci
296. Saprolegnia diclina vs20
297. Saprolegnia parasitica cbs 223 65
298. Pythium iwayamai
299. Pythium arrhenomanes
300. Pythium ultimum
301. Pythium aphanidermatum
302. Pythium irregulare
303. Pythium vexans
304. Hyaloperonospora arabidopsidis
305. Phytophthora kernoviae
306. Phytophthora ramorum
307. Phytophthora lateralis
308. Phytophthora sojae
309. Phytophthora parasitica
310. Phytophthora infestans
311. Thalassiosira oceanica
312. Thalassiosira pseudonana
313. Phaeodactylum tricornutum
314. Galdieria sulphuraria
315. Chondrus crispus
316. Nitrosopumilus maritimus scm1
317. Caldisphaera lagunensis dsm 15908
318. Metallosphaera cuprina ar 4
319. Metallosphaera sedula
320. Sulfolobus tokodaii str 7
321. Sulfolobus islandicus hve10 4
322. Sulfolobus solfataricus 98 2
323. Sulfolobus acidocaldarius dsm 639
324. Hyperthermus butylicus dsm 5456
325. Thermogladius cellulolyticus 1633
326. Ignicoccus hospitalis kin4 i
327. Staphylothermus hellenicus dsm 12710
328. Staphylothermus marinus f1
329. Desulfurococcus kamchatkensis 1221n
330. Desulfurococcus mucosus dsm 2162
331. Thermofilum pendens hrk 5
332. Aciduliprofundum boonei t469 gca 000025665 1
333. Methanosaeta thermophila pt
334. Methanothermobacter marburgensis str marburg
335. Methanobrevibacter ruminantium m1
336. Methanosphaera stadtmanae dsm 3091
337. Methanothermus fervidus dsm 2088

## Appendix B: Ranked top 50 genes for each BLAST E-value threshold

### Ranked list, Top50 (TM-containing, ER-localized only), BLAST threshold: 0

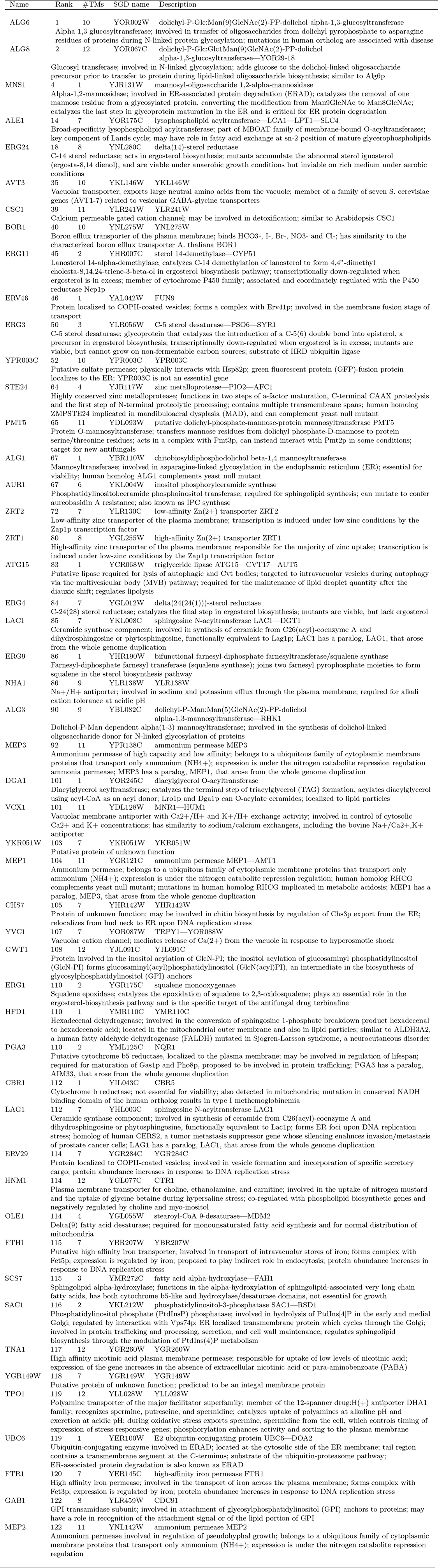

### Ranked list, Top50 (TM-containing, ER-localized only), BLAST threshold: 1e-2

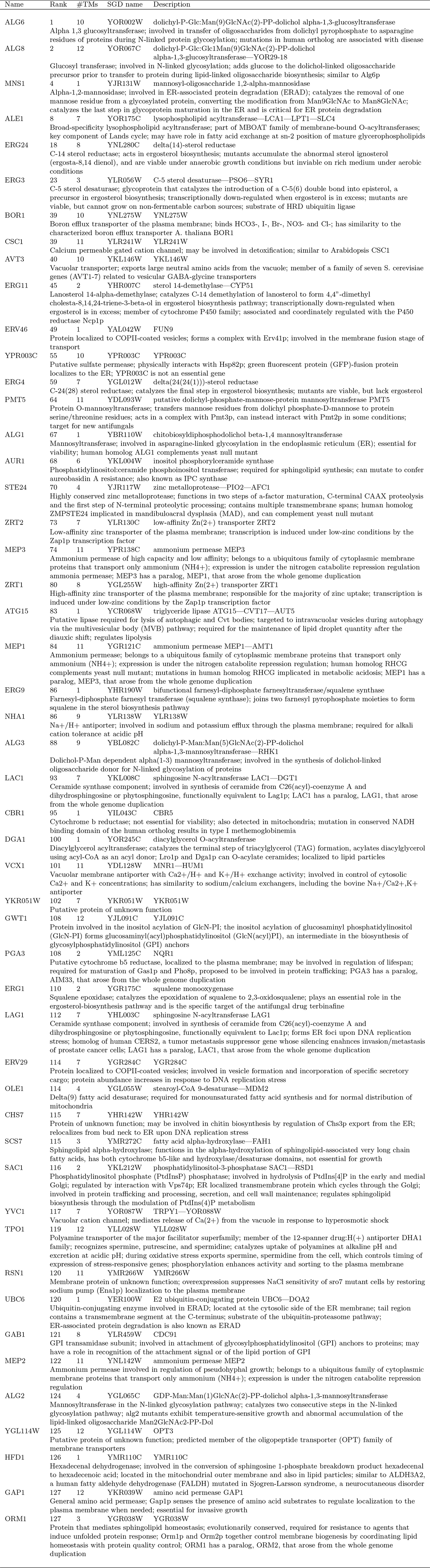

### Ranked list, Top50 (TM-containing, ER-localized only), BLAST threshold: 1e-10

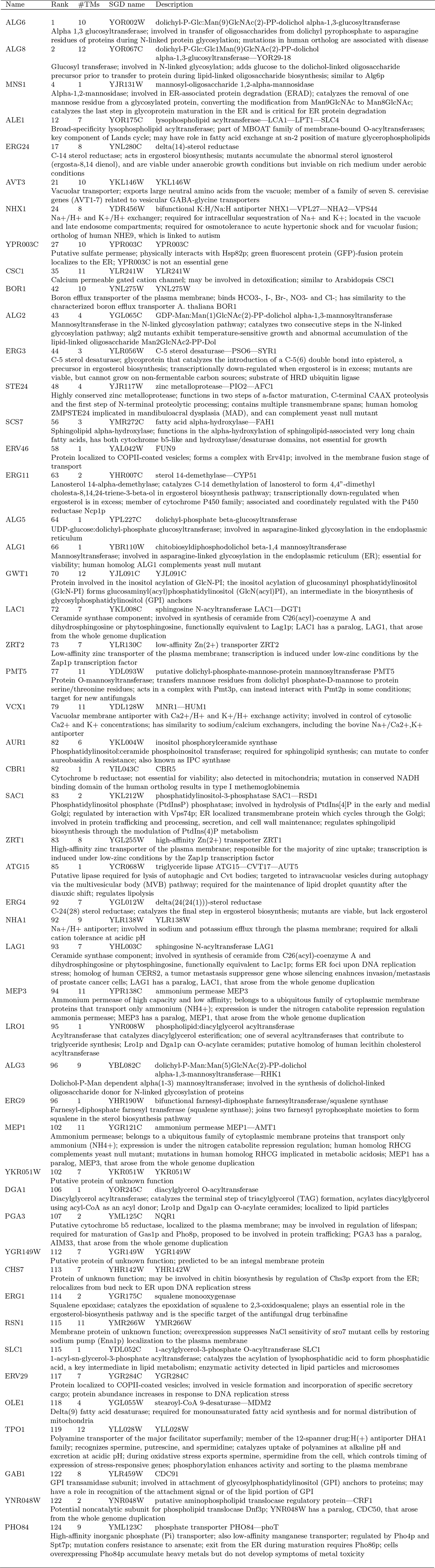

